## Supplementary material for "Modular assembly of dynamic models in systems biology": S1 Text

Supplementary Information

Michael Pan, Peter J. Gawthrop, Joseph Cursons and
Edmund J. Crampin

---

**B A white-box modularity approach to the MAPK cascade** **6**

### A Parameter inference

#### A.1 Alternative formulations of thermodynamic parameters

Depending on the available data, one can write species and reaction equations such that the parameters have dimension energy. The equation for the species (Eq. 3 in the Main Text) could be written as

$$\mu = \mu^0 + RT\ln(x/x^0) \tag{S1}$$

where $\mu^0$ is the standard chemical potential taken at a concentration of $x^0$ (for simplicity, $x^0$ is often taken to be 1M). The thermodynamic parameters are related by $K = (1/x^0)\exp(\mu^0/RT)$. Similarly, the equation corresponding to the reactions (Eq. 4 in the Main Text) could also be written as

$$v = \kappa^* \exp\left(\frac{E^*}{RT}\right)\left[\exp\left(\frac{A^f}{RT}\right) - \exp\left(\frac{A^r}{RT}\right)\right] \tag{S2}$$

where $E^* = RT\ln(\kappa/\kappa^*)$ and $\kappa^* = 1$M/s is used for unit consistency. The $E^*$ parameter is linked to the activation energy of the reaction.

#### A.2 MAPK model

The parameters of the core MAPK model were chosen to match the behaviour of the Huang and Ferrell [1] model as closely as possible. In general, there are families of parameter sets with identical kinetic behaviour, so there are an infinite number of plausible thermodynamic parameters. To obtain a unique solution here, we assume that $K_{\text{MAP4K}} = K_{\text{MAP3K}} = K_{\text{MAP2K}} = K_{\text{MAPK}} = K_{\text{MAP3K-Pase}} = K_{\text{MAP2K-Pase}} = K_{\text{MK-Pase}} = 1\ \mu\text{M}^{-1}$. We additionally assume that $K_{\text{ATP,hyd}} = e^{-\Delta G_{\text{ATP}}/RT}\ \mu\text{M}^{-1}$, $K_{\text{ADP}} = 1\ \mu\text{M}^{-1}$ and $K_{\text{Pi}} = 1\ \mu\text{M}^{-1}$ to fit a physiological free energy of ATP hydrolysis of $\Delta G_{\text{ATP}} = -50$ kJ/mol. While parameter uncertainty is not dealt with in this manuscript, the set of plausible parameters can be formalised in terms of null spaces of the stoichiometric matrix, and we refer readers to other papers [2–4] for further information on parameter uncertainty.

The primary motif that we consider for parameter estimation will be the phosphorylation cycle

$$X + E \underset{d_E}{\overset{a_E}{\rightleftharpoons}} XE \underset{l_E}{\overset{k_E}{\rightleftharpoons}} XP + E \tag{S3a}$$

$$XP + F \underset{d_F}{\overset{a_F}{\rightleftharpoons}} XF \underset{l_F}{\overset{k_F}{\rightleftharpoons}} X + F \tag{S3b}$$

where X is the dephosphorylated substrate, XP is the phosphorylated substrate, E is the kinase and F is the phosphatase. As with Huang and Ferrell, we have omitted ATP, ADP and Pi from the reaction scheme; in reality, the kinetic parameters are dependent on their concentrations.

The kinetic parameters are identical between phosphorylation cycles; $a_E = a_F = a = 1000\ \mu\text{M}^{-1}\text{s}^{-1}$, $d_E = d_F = d = 150\ \text{s}^{-1}$ and $k_E = k_F = k = 150\ \text{s}^{-1}$. Huang and Ferrell also assume that $l_E = l_F = 0$. However, this is not possible in reality because all reactions are reversible. Indeed, their values are constrained by the laws of thermodynamics. Specifically, because all four reactions form a cycle that consumes ATP and produces ADP and Pi with a free energy of $\Delta G_{\text{ATP,hyd}}$, detailed balance states that

$$\frac{a_E k_E a_F k_F}{d_E l_E d_F l_F} = e^{-\Delta G_{\text{ATP,hyd}}/RT} \tag{S4}$$

and hence

$$l_E l_F = \frac{a^2 k^2}{d^2} e^{\Delta G_{\text{ATP,hyd}}/RT} \tag{S5}$$

In other words, the more favourable ATP hydrolysis is, the smaller $l_E$ and $l_F$ become and the closer they are to the ideal state of $l_E = l_F = 0$. However, since there is a finite amount of energy from ATP hydrolysis in reality, the values of $l_E$ and $l_F$ are positive.

We minimise the magnitudes of $l_E$ and $l_F$ by minimising the objective function

$$J = l_E^2 + l_F^2 = l_E^2 + D^2/l_E^2 \tag{S6}$$

where $D = (a^2k^2)/(d^2)e^{\Delta G_{\text{ATP}}/RT}$. The minimum occurs at

$$dJ/dl_E = 2l_E - 2D^2/l_E^3 = 0 \tag{S7}$$

and solving for $l_E$, we find that

$$l_E = l_F = D^{1/2} \tag{S8}$$

By relating kinetic to thermodynamic parameters, the thermodynamic parameters can be calculated using the following equations:

$$K_{\text{XE}} = K_{\text{X}} K_{\text{E}} (d/a) e^{\mu_{\text{ATP}}/RT} \tag{S9a}$$
$$\kappa_{\text{E1}} = d/K_{\text{XE}} \tag{S9b}$$
$$\kappa_{\text{E2}} = k/K_{\text{XE}} \tag{S9c}$$
$$K_{\text{XP}} = K_X D (d/ak) e^{(\mu_{\text{ATP}} - \mu_{\text{ADP}})/RT} \tag{S9d}$$
$$K_{\text{XF}} = K_{\text{XP}} K_{\text{F}} (d/a) \tag{S9e}$$
$$\kappa_{\text{F1}} = d/K_{\text{XF}} \tag{S9f}$$
$$\kappa_{\text{F2}} = k/K_{\text{XF}} \tag{S9g}$$
$$\tag{S9h}$$

where $\kappa_{ij}$ is the rate parameter for reaction $j$ of enzyme $i$ and we set $T = 310$ K.

With the value of $K_{\text{XP}}$ determined, this procedure can be applied to the next cycle along the pathway and propagated throuout the entire cascade to find all thermodynamic parameters.

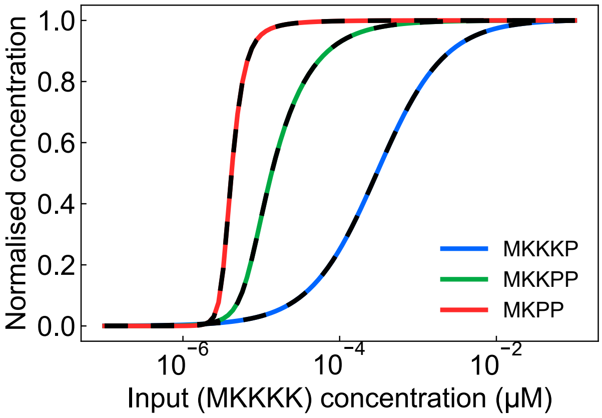

**Figure S1: Comparison of the bond graph model of the MAPK cascade to the Huang and Ferrell model.** The solid lines are identical to those in Figure 5B. The dotted lines are generated from running the Huang and Ferrell model under the same initial conditions.

A comparison of the bond graph model with the kinetic model is shown in Figure S1. While the two models differ in the reversibility of the catalytic reaction, these differences turn out to be negligible under physiological values of $\Delta G_{\mathrm{ATP,hyd}}$.

The same method was used to determine parameters for the feedback loops, but the $\kappa$ parameters for the phosphatase enzymes were scaled up by a factor of 10. This was to allow the feedback to occur within the range of input concentrations where ultrasensitivity is typically observed.

#### A.3 Glycolysis model

The models of glycolysis were based on the *E. coli* model developed by Mason and Covert [2]. The generalised kinetics model used parameters directly from Mason and Covert, with parameters converted using the methods in § A.1. Out of the multiple parameterisations of the model, we chose the one with the shortest response time (bottom row of Fig. 5 of their paper, with penalty weight of 100), which appeared to be the most realistic given that the typical response time for glycolysis is on the order of seconds [5]. The dilution effects of growth were assumed to be negligible.

We then chose parameters for the mass action and Michaelis-Menten models to match important behaviours of the more complex generalised kinetics model. Since the species components are identical between all models, their thermodynamic parameters were unchanged between the models.

For the mass action model, we chose the reaction parameters $\kappa$ such that the fluxes at the reference steady state would be identical to the generalised kinetics model. Specifically, the reaction parameter $\kappa$ for each reaction was determined by rearranging Eq. 4 of the Main Text:

$$\kappa = v \left[ \exp\left(\frac{A^f}{RT}\right) - \exp\left(\frac{A^r}{RT}\right)\right]^{-1} \tag{S10}$$

where $v$ was obtained by simulating the generalised kinetics model to steady state and the reaction affinities $A^f$ and $A^r$ was inferred from the chemical potentials of each species.

For the Michaelis-Menten model, we chose parameters to match the dynamic behaviour of the system under reference chemostat concentrations. The Michaelis-Menten equation contains only two binding parameters: one for the forward complex and one for the reverse complex. We chose these binding parameters to match the binding properties of the carbon species. For all reactions apart from *fba* (which we will deal with later as a separate case) there is only a single carbon species in the substrates and products; we denote these by $s^*$ and $p^*$ respectively. Then, assuming the concentrations of the other external species are constant, we can rewrite the generalised kinetics equation (Eq. 12 of the Main Text) as

$$v = \bar{\kappa}_{\text{GK}} e_0 \frac{e^{A^f/RT} - e^{A^r/RT}}{-1 + \prod_{s\in\mathcal{S}}\left(1 + \frac{e^{\mu_s/RT}}{R_{b,s}}\right) + \prod_{p\in\mathcal{P}}\left(1 + \frac{e^{\mu_p/RT}}{R_{b,p}}\right)} \tag{S11a}$$

$$= \bar{\kappa}_{\text{GK}} e_0 \frac{e^{A^f/RT} - e^{A^r/RT}}{-1 + \left(1 + \frac{e^{\mu_{s^*}/RT}}{R_{b,s^*}}\right) C_f + \left(1 + \frac{e^{\mu_{p^*}/RT}}{R_{b,p^*}}\right) C_r} \tag{S11b}$$

$$= \bar{\kappa}_{\text{GK}} e_0 \frac{e^{A^f/RT} - e^{A^r/RT}}{(C_f + C_r - 1) + \frac{e^{\mu_{s^*}/RT}}{R_{b,s^*}} C_s + \frac{e^{\mu_{p^*}/RT}}{R_{b,p^*}} C_r} \tag{S11c}$$

where

$$C_f = \prod_{s\in\mathcal{S}\backslash\{s^*\}} \left(1 + \frac{e^{\mu_s/RT}}{R_{b,s}}\right) \tag{S12a}$$

$$C_r = \prod_{p\in\mathcal{P}\backslash\{p^*\}} \left(1 + \frac{e^{\mu_p/RT}}{R_{b,p}}\right) \tag{S12b}$$

Additionally, we can define

$$A^f_{\text{side}} = \sum_{s\in\mathcal{S}\backslash\{s^*\}} \mu_s = A^f - \mu_{s^*} \tag{S13a}$$

$$A^r_{\text{side}} = \sum_{p\in\mathcal{P}\backslash\{p^*\}} \mu_p = A^r - \mu_{p^*} \tag{S13b}$$

which allows us to rewrite the generalised kinetics equation in the form of the Michaelis-Menten equation.

$$v = \frac{\bar{\kappa}_{\text{GK}}}{C_f + C_r - 1} e_0 \frac{e^{A^f/RT} - e^{A^r/RT}}{1 + \frac{e^{A^f/RT} e^{-A^f_{\text{side}}/RT}}{R_{b,s^*}(C_f + C_r - 1)} C_f + \frac{e^{A^r/RT} e^{-A^r_{\text{side}}/RT}}{R_{b,p^*}(C_f + C_r - 1)} C_r} \tag{S14}$$

By comparing constants to Eq. 11 of the Main Text, we can determine the parameters for Michaelis-Menten kinetics under reference chemostat concentrations.

$$\bar{\kappa}_{\mathrm{MM}} = \frac{\bar{\kappa}_{\mathrm{GK}}}{C_f + C_s - 1} \tag{S15a}$$

$$R_{b0} = R_{b,s^*} \frac{(C_f + C_r - 1)e^{A^f_{\mathrm{side}}/RT}}{C_f} \tag{S15b}$$

$$R_{b1} = R_{b,p^*} \frac{(C_f + C_r - 1)e^{A^r_{\mathrm{side}}/RT}}{C_r} \tag{S15c}$$

For the reaction *fba*, the above approach is not possible as there are two carbon products. The generalised kinetics and Michaelis-Menten rate laws for *fba* can be written as

$$v_{\mathrm{fba,GK}} = \bar{\kappa}_{\mathrm{GK}} e_0 \frac{e^{\mu_{\mathrm{F16P}}/RT} - e^{(\mu_{\mathrm{DHAP}}+\mu_{\mathrm{GAP}})/RT}}{1 + \frac{e^{\mu_{\mathrm{F16P}}/RT}}{R_{b,\mathrm{F16P}}} + \frac{e^{\mu_{\mathrm{DHAP}}/RT}}{R_{b,\mathrm{DHAP}}} + \frac{e^{\mu_{\mathrm{GAP}}/RT}}{R_{b,\mathrm{GAP}}} + \frac{e^{(\mu_{\mathrm{DHAP}}+\mu_{\mathrm{GAP}})/RT}}{R_{b,\mathrm{DHAP}}R_{b,\mathrm{GAP}}}} \tag{S16a}$$

$$v_{\mathrm{fba,MM}} = \bar{\kappa}_{\mathrm{MM}} e_0 \frac{e^{\mu_{\mathrm{F16P}}/RT} - e^{(\mu_{\mathrm{DHAP}}+\mu_{\mathrm{GAP}})/RT}}{1 + \frac{e^{\mu_{\mathrm{F16P}}/RT}}{R_{b0}} + \frac{e^{(\mu_{\mathrm{DHAP}}+\mu_{\mathrm{GAP}})/RT}}{R_{b1}}} \tag{S16b}$$

Thus the two rate laws only differ functionally only in the presence of the binding terms for DHAP and GAP in the denominator. Because the two rate laws are identical when the terms involving $\mu_{\mathrm{DHAP}}$ and $\mu_{\mathrm{GAP}}$ are neglected, we set $\bar{\kappa}_{\mathrm{MM}} = \bar{\kappa}_{\mathrm{GK}}$ and $R_{b0} = R_{b,\mathrm{F16P}}$. The final parameter $R_{b1}$ was chosen to match $v_{\mathrm{fba,ss}}$, the flux of the reaction at steady state under reference conditions:

$$R_{b1} = \frac{e^{(\mu_{\mathrm{DHAP}}+\mu_{\mathrm{GAP}})/RT}}{\frac{\bar{\kappa}_{\mathrm{MM}} e_0}{v_{\mathrm{fba,ss}}} \left[e^{\mu_{\mathrm{F16P}}/RT} - e^{(\mu_{\mathrm{DHAP}}+\mu_{\mathrm{GAP}})/RT}\right] - \frac{e^{\mu_{\mathrm{F16P}}/RT}}{R_{b0}} - 1} \tag{S17}$$

#### B A white-box modularity approach to the MAPK cascade

In the traditional bond graph approach to hierarchical modelling, modules are specified through the use of fixed ports that are unable to be changed after the development of a model. However, it is increasingly acknowledged that white-box modularity is important to constructing large-scale models in systems biology. This approach requires both the internal details of a model to be externally accessible and for connections between components to be modified as needed. Here we illustrate that bond graphs are compatible with white-box modularity by defining the MAPK cascade model in terms of white-box modules.

To help understand white-box modularity in the context of bond graphs, we first consider the merging of kinase and phosphatase models into a model of a phosphorylation cycle (Figure S2A–C). In a white-box approach, the kinase and phosphatase modules are specified as closed systems with no external connections

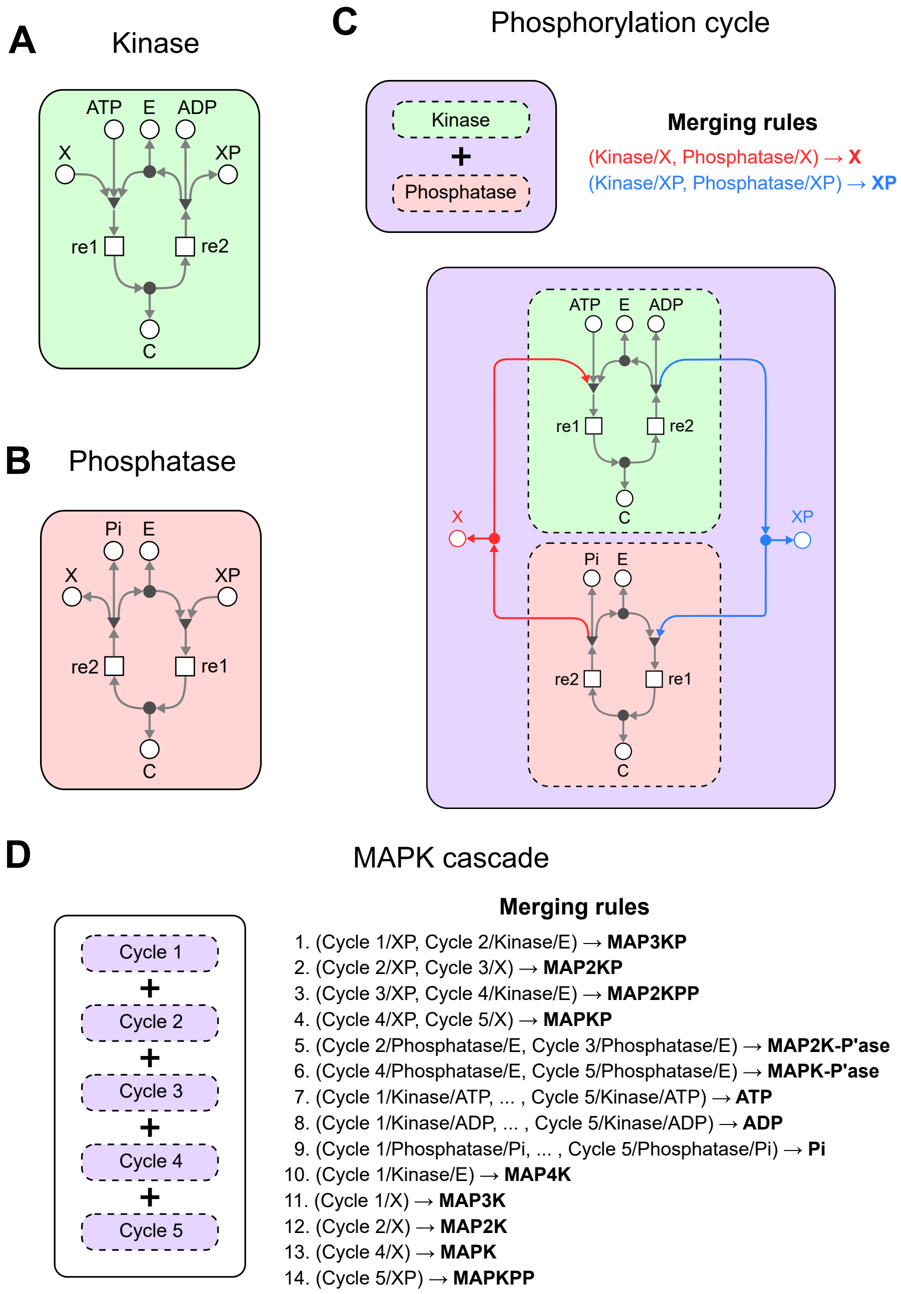

**Figure S2: A white-box approach to defining a model of the MAPK cascade.** Similar to the black-box approach, modules are defined for the **(A)** kinase and **(B)** phosphatase. However, unlike the black-box approach, individual modules are themselves independent models that are simulatable. **(C)** The white-box modularity approach is illustrated on the phosphorylation cycle. (Top) Large models are composed from smaller models through a set of "merging rules" that link components representing the same biological entity. Here, the two merging rules correspond to the dephosphorylated substrate X (red) and the phosphorylated substrate XP (blue). (Bottom) Components are merged by disconnecting the components from each respective module, exposing the respective port and connecting them to the same component through a conservation of mass relation; the modified connections are colour-coded in red and blue for X and XP respectively. **(D)** A model of the MAPK cascade can be constructed by composing multiple copies of the phosphorylation cycle in (C), with the merging rules shown on the right.

(Figure S2A–B). The coupled phosphorylation loop model can then be defined as the composition of the constituent kinase and phosphatase models, together with a set of "merging rules" that define the components that are shared between the two models (Figure S2, top panel). Here we denote the unification of $n$ components using the notation

$$(\text{Component 1}, \text{Component 2}, \ldots, \text{Component } n) \rightarrow \textbf{Merged component} \tag{S18}$$

When two or more components within different models are merged together, the original components are disconnected and removed from their original models, being replaced by external ports. These ports are then connected to a new shared component through a mass conservation law. We use forward slashes to represent the hierarchical structure of the components within modules; for example, "Kinase/X" refers to the X component within the Kinase module. Therefore, in the case of the phosphorylation loop, the shared components correspond to the unphosphorylated substrate X and phosphorylated substrate XP that occur in both the kinase and phosphatase modules (Figure S2C). The red and blue colour code indicates the components and connections that are modified to merge the components together.

When two or more non-identical components are merged together, the modeller must decide which of the original components to use. This is an important feature of the bond graph approach; inconsistencies between models are flagged to the modeller to act on. To simplify the analysis of this example, we assume that the components being merged are identical. Dealing with parameter inconsistencies between models is the subject of future work.

Using this notion of model composition, the full model of the MAPK pathway in Figure 4D could equivalently be defined using the white-box definition in Figure S2D. The merging rules are grouped as follows:

- Rules 1–6 connect identical species between each of the phosphorylation loops
- Rules 7–9 merge the sources of ATP, ADP and Pi used in each of the loops
- Rules 10–14 promote some species to the highest level of the model hierarchy for convenience

As with other approaches to white-box modularity, this approach provides the modeller with additional flexibility, since models can be individually simulated at each level of the model hierarchy. From the perspective of software engineering, attributes can be assigned to components at each level of the modelling hierarchy. This could help allow a submodel's parameters to be inherited by a parent model, or to document a change to the parameter of a model during the merging process [6]. While the merging rules were manually specified here, it is envisioned that the components corresponding to each species could be labelled using annotations so that models can be merged automatically.
